# Minimally invasive, pressure probe based sampling allows for *in-situ* gene expression analyses in plant cells

**DOI:** 10.1101/768978

**Authors:** Hiroshi Wada, Simone D. Castellarin, Mark A. Matthews, Kenneth A. Shackel, Gregory A. Gambetta

## Abstract

**Background:** Gene expression analyses are conducted using multiple approaches and increasingly research has been focused on assessing gene expression at the level of a tissue or even single-cells. To date, methods to assess gene expression at the single-cell in plant tissues have been semi-quantitative, require tissue disruption, and/or involve laborious, possibly artifact-inducing manipulation. In this work, we used grape berries (*Vitis vinifera* L. Zinfandel) as a model in order to examine the validity and reproducibility of an *in-situ* gene expression analysis method combining a cell pressure probe (CPP) with quantitative PCR (qPCR).

**Results:** We developed a method to directly assess gene expression levels via qPCR from cellular fluids sampled *in-situ* with a CPP. Cellular fluids, with volumes in the picoliter range, were collected from intact berries with a CPP at various depths across skin and mesocarp tissues. The expression of a key anthocyanin biosynthetic gene, UDP-glucose: flavonoid 3-O-glucosyltransferase (*VviUFGT*), was analyzed as a test case since its expression is restricted to cells producing anthocyanins in grape berry skins during ripening. The method identifies samples contaminated with significant levels of genomic DNA by amplifying a region of *VviUFGT* that spans an intron. Therefore false positives were discarded which occurred in 28% of the samples tested. Shallow probing of skin cells showed high *VviUFGT* expression as expected while deeper probing of mesocarp cells resulted in no *VviUFGT* expression.

**Conclusions:** The clear correspondence of *VviUFGT* expression to the targeted cell samples suggests that the *in-situ* gene expression analysis using a CPP is reliable and does not result in contamination as the probe moves through tissues. This method can be paired to single-cell transcriptomic analyses in the future. We conclude that this technique represents a minimally invasive method of sampling plant cells *in-situ* which creates an opportunity for the analysis of cellular level, spatiotemporal responses in heterogeneous plant tissues.

## Background

Single cell analyses have been conducted at the transcript and metabolite levels over the last two decades, and some have succeeded in the characterization of molecular information in individual plant cells (Kehr 2003; Efroni and Birnbaum 2016; Yuan et al. 2017). Nevertheless these analyses are extremely technically challenging and are semi-quantitative, require tissue disruption, and/or involve laborious, possibly artifact-inducing manipulation. For gene expression analyses, *in-situ* hybridization has been used fairly extensively (e.g., Drea et al. 2009). This method visualizes the spatial distribution of target gene expression in fixed and sectioned tissues via microscopy, but information is limited to the presence or absence of expression. Laser capture microdissection (LCM) is powerful, but still requires sectioning (most commonly via cryo-dissection) which may be challenging for certain tissues such as fruit cells. During LCM, whether the cell components in samples might be altered when the tissues are subjected to heat and radiation from an infrared laser remains questionable (Kehr 2003), implying the need of confirmation using some direct approach. Fluorescent labeled sorting methods have also been utilized, which requires tissue digestion after dissection (Yuan et al. 2017) raising the possibility that digested samples may lose cell-specific information. Therefore, a direct *in-situ* method that allows cell-specific analysis could prove extremely valuable.

The cell pressure probe (CPP) (Hüsken et al. 1978) has traditionally been used for directly determining the physicochemical parameters of individual plant cells, such as cell turgor and hydraulic conductivity, by inserting a fine oil-filled microcapillary tip into the cell, and monitoring/adjusting the location of oil/sap boundary (meniscus) under a microscope (Steudle 1993; Tomos and Leigh 1999). This minimally invasive approach has been used to quantify cell-to-cell variation in water relations *in-situ* (Nonami and Schulze 1989). The CPP has also been used to collect cellular fluids for metabolite analyses in order to reveal spatial variations in cellular metabolism using enzymatic assays (Koroleva et al. 1997) and more recently mass spectrometry-assisted cell metabolomics (Nakashima et al. 2016; Wada et al. 2019).

Cellular level gene expression assays have been successful using RT-PCR on cellular fluids obtained either with the CPP (Brandt et al. 1999, 2002; Gallagher et al. 2001; Schliep et al. 2010) or simple aspiration using a glass microcapillary (Jones and Grierson 2003). These previous studies acknowledged that when collecting the cellular fluids using glass microcapillaries it is possible that the sample would be contaminated with genomic DNA (gDNA) which can lead to artifacts during the PCR amplification (Brandt 2005).

To date, no attempt has been made to quantify gene expression at the single cell level using the CPP and quantitative PCR (qPCR) while simultaneously assessing levels of gDNA contamination. In this work, we have developed a protocol coupling single cell sampling with the CPP to qPCR analyses along with primer designs that identify contaminating gDNA. By using intact ripening grape berries that accumulated anthocyanins in the skin cells, we tested the validity of this protocol for the *in-situ* quantification of gene expression in intact grape berry cells.

## Results and Discussion

### Identifying gDNA contamination

The expression of a key anthocyanin biosynthetic gene, UDP-glucose: flavonoid 3-O-glucosyltransferase (*VviUFGT*) was analyzed as a test case since its expression is restricted to cells producing anthocyanins in grape berry skins during ripening. In order to identify instances of gDNA contamination of the collected cellular fluids we targeted a region of *VviUFGT* that spanned an intron sequence (**Fig. 1**). After the qPCR was carried out all reactions were visualized by gel electrophoresis allowing the identification of gDNA contamination that occurred in 28.6% (6/21) of the samples.

**Fig. 1.**
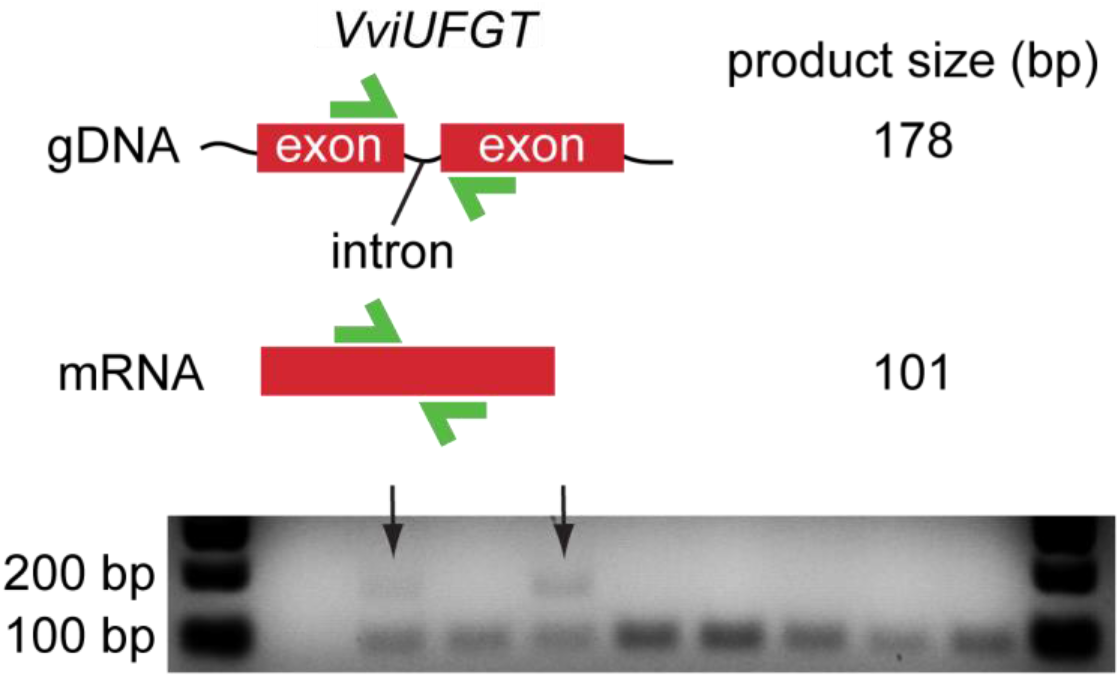
The identification of gDNA contamination of the collected cellular fluids. A region of *VviUFGT* was targeted for qPCR analyses that spanned an intron sequence such that the resulting product from gDNA template was significantly larger that the desired product from a mRNA. Following qPCR all reactions were visualized by gel electrophoresis allowing the identification of gDNA contamination (black arrows).

As cell contents are collected with the CPP there is some chance that the tip will pass through and/or insert into nuclei which would result in gDNA contamination. This issue is critically important if sensitive quantitative methods such as qPCR will be used downstream, and has received no attention in some studies (Laval et al. 2002; Schliep et al. 2010). Brandt et al. (1999) reported that their controls for gDNA contimination, “produced no detectable signals in nearly all experiments”, suggesting that the incidence of gDNA contaminaion in their work was extremely low. In the curent study the rate was significant, reinforcing the need to include a robust method of identification of gDNA contamination in *in-situ* cell sampling methodologies.

Differences in the rates of gDNA contamination may result because of differences in the tissue from which cells are sampled. In ripening grape berries the diameter of nuclei in subepidermal cells are 2.5-5 μm (Diakou and Carde 2001), similar to the tip inner diameter we used here (see Methods). Assuming that the nucleus diameters were similar in all cell types across tissues, it is possible that the tip might have poked the nucleus in the target cells below epidermis. In this context, the extent of gDNA contamination would depend on the size and localization of nucleus (nuclei) in the cytosol, as well as the volumetric ratio of cytosolic space to cell volume. Additionally, as the depth of probing increases so would the likelihood of gDNA contamination since the CPP would have to pass through more cell layers prior to sampling. In the current study, of the 6 samples that were positive for gDNA contamination, 4 of the samples were from probing depths of >1000 μm.

### *In-situ* CPP sampling and qPCR

In this study, cellular fluids from individual grape berries were extracted using the CPP and the cell sap was used directly in downstream qPCR analyses (**Fig. 2**). Different probing techniques were used in order to test different tissues (**Fig. 3**). Shallow and deep probing techniques were intended to extract cell fluids from the skin and mesocarp tissues respectively. The continuous probing method extracted cell fluids from numerous cells as the probe moved deeper allowing for a greater volume of cell fluids as a control.

**Fig. 2.**
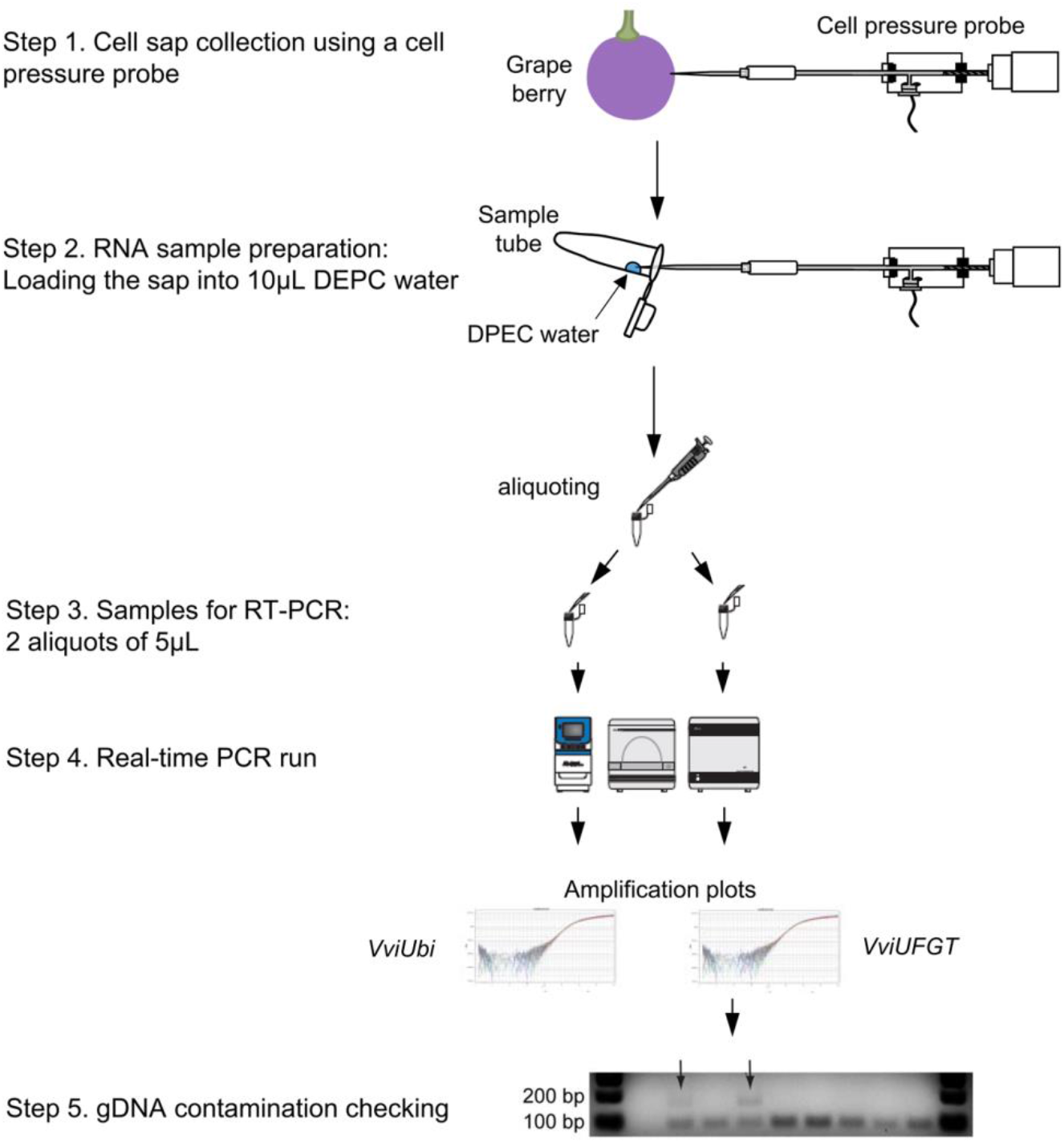
Illustrated workflow of in-situ cellular gene expression in post-veraison grape berries.

**Fig. 3.**
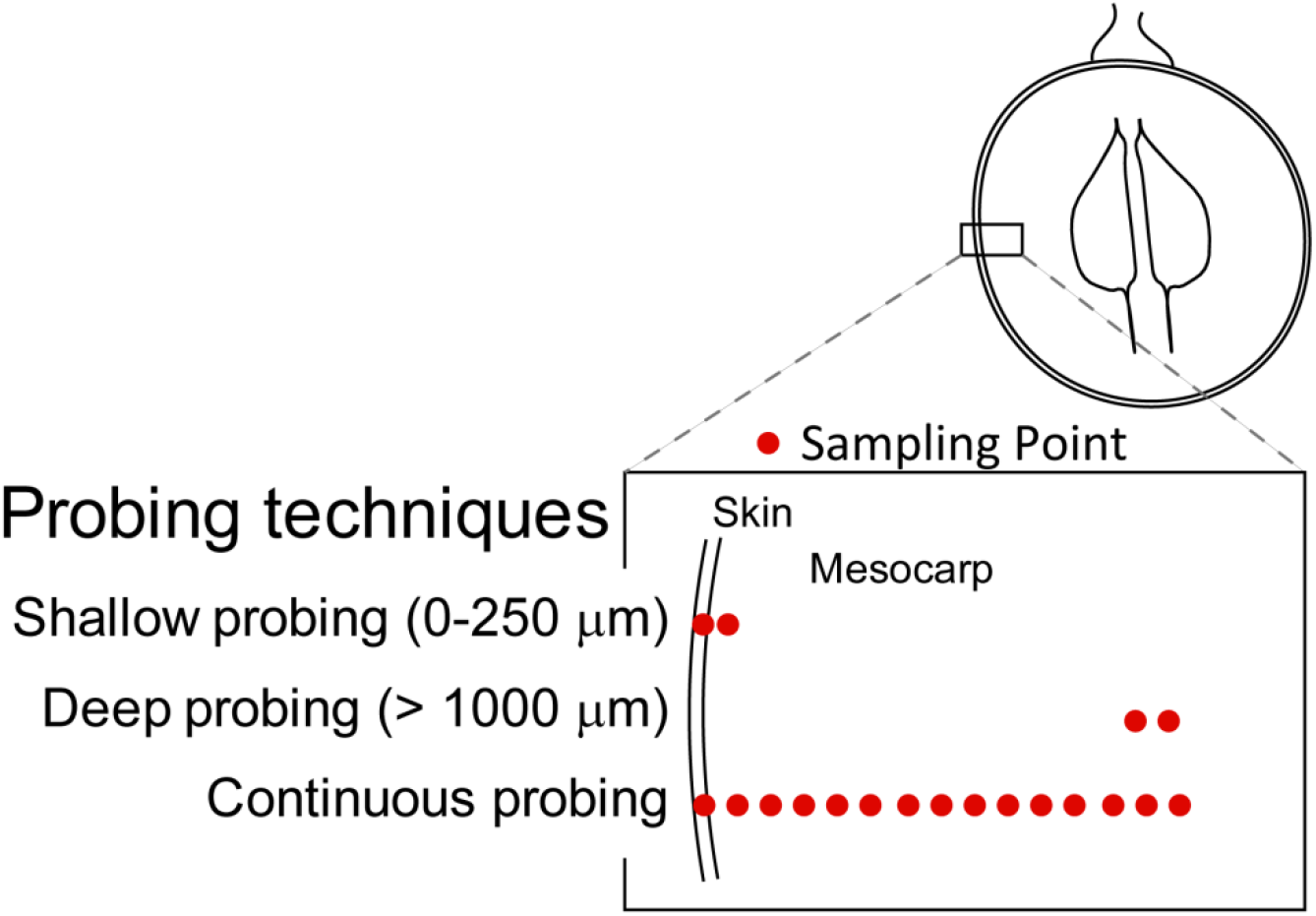
Probing techniques used in this study. The cell layers collected in three extraction methods, shallow, deep, and continuous probing were shown in red.

In total 34 samples were collected with the CPP. The extensively utilized grapevine housekeeping gene Ubiquitin1 (*VviUbi*) was used as a positive control and reference gene (Bogs et al. 2005; Castellarin et al. 2007b, a, 2011) and of those 34 samples 21 (~62%) exhibited *VviUbi* expression. There was a clear difference in the *VviUFGT* gene expression corresponding to the different probing techniques (**Fig. 4**) which corresponded to different specific depths within the berry, and thus different tissues. Shallow and continuous probing included skin cells and exhibited *VviUFGT* expression at high frequencies (>80%). Contrastingly, there was no *VviUFGT* expression in mesocarp cells greater than 250μm below epidermis, corresponding to cells with no coloration (**Fig. 4**). Of those samples exhibiting *VviUFGT* expression the average expression, relative to *VviUbi*, was 0.61 ± 0.15 (standard error) which is extremely similar to previous studies using the same reference gene in Cabernet Sauvignon (Castellarin et al. 2007a).

**Fig. 4.**
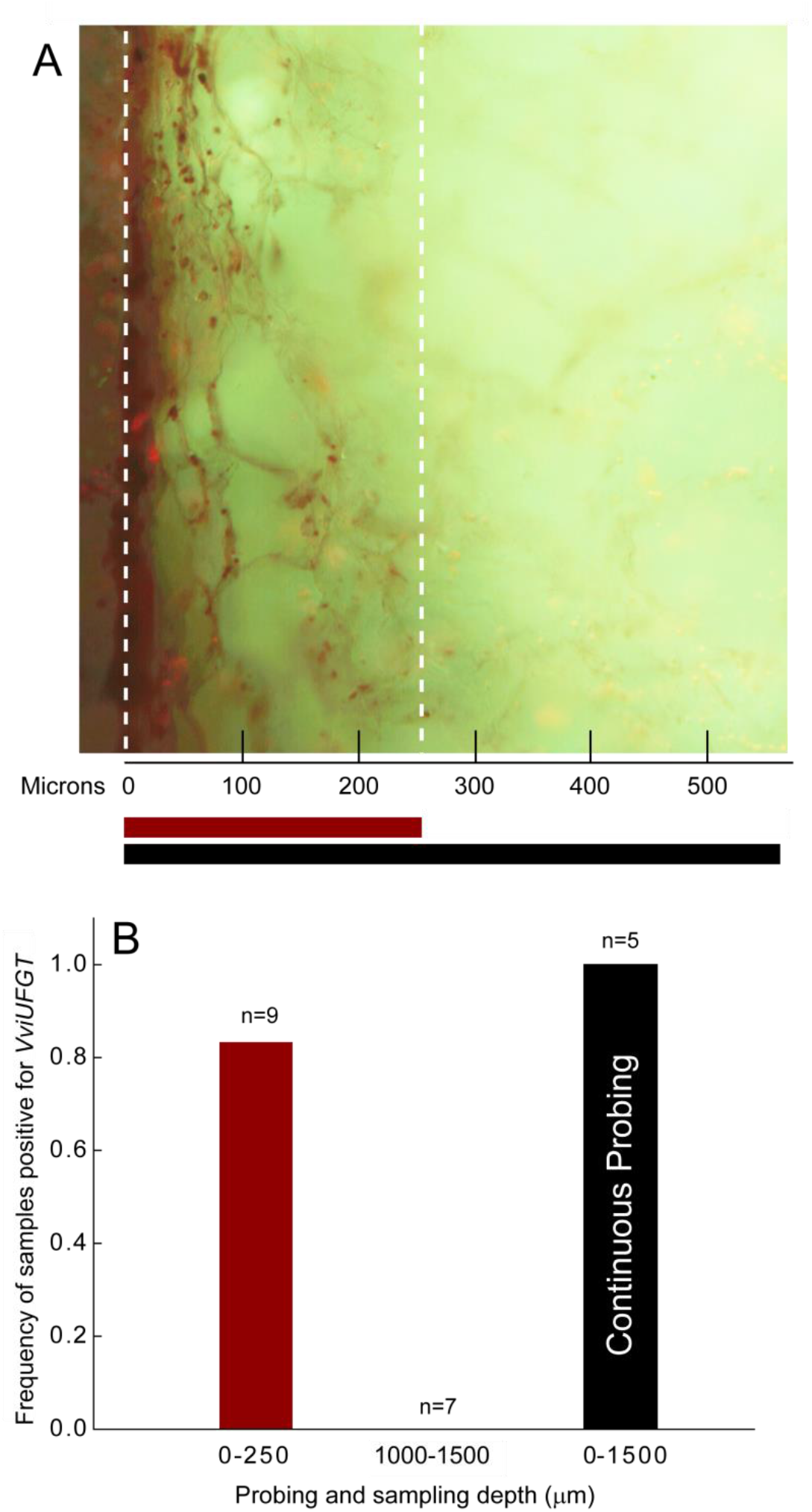
Analysis of *VviUFGT* expression across sampled tissues. (a), image of cross section of grape berry. Red and black bars in A correspond to the shallow and continuous probing. (b), frquency of samples that tested positive for *VviUFGT* expressions in cell sap colllected by performing shallow depth (0-250 μm), deeper depth (1000-1500 μm), and continuous probing techniques in postveraison berries. Zero samples were positive for *VviUFGT* expression at the 1000-1500 μm depth.

In this work the overall success ratio of the gene expression assay was 62%, which was similar to Jones and Grierson (2003) (64% in *Arabidopsis thaliana* root hair cells). It has been suggested that the use of cellular gene expression methods in plant cells is more difficult than in animal cells (Brandt 2005) because plant cells have rigid cell walls and small cytosolic spaces due to the presence of large vacuoles. Therefore a sampling success rate of >60% is reasonable, given the technical challenges involved, and is feasible in terms of experimentation.

Most previous gene expression analyses using CPP were only semi-quantitative (Brandt, 2005). In the current study we demonstrate a minimally invasive *in-situ* method that produces a reliable quantification of target gene expression in a single qPCR run while simultaneously assessing the presence of gDNA contamination. Although it is not possible to visually identify the cell being sampled, the use of a Piezo-manipulator for precise depth control, and appropriate manipulation of oil pressure, in the current study increases the precision of sampling cells from targeted tissues. However, in extremely complex tissues this could be a limitation.

## Conclusions

We combined qPCR analysis with CPP sampling to establish a new protocol for *in-situ* cellular gene expression analysis in plant cells. Utilizing qPCR primers designed to detect an intron sequence successfully identified instances of gDNA contamination. The gene expression analyses demonstrated the reliability and robustness of the method which was able to quantify expression of our test gene from as few as two sampled cells. Prior knowledge of spatial turgor distribution and hence appropriate pressure manipulation are required to reduce the possible risk of contamination from the non-target cells. Further improvement of both the resolution and sensitivity of detection will be required for performing more global gene expression analyses in picolitre sap samples. This technique represents a minimally invasive method of sampling plant cells *in-situ*, and could potentially contribute to a better understanding of cell heterogeneity in plant tissues.

## Methods

### Plant material

Grape berries (*Vitis vinifera* L. ‘Zinfandel’) were sampled from field-grown vines located at the University of California, Davis, CA, USA (38.32’ N latitude and 121.46’ W longitude, elevation 18 m above sea level) according to Castellarin et al. (2016). The anthesis date was the day on which 50% of the cluster was flowering, and developmental time is represented as days after anthesis (DAA). Berry skins began to accumulate anthocyanins at 69 DAA. Ripening berries were randomly collected at 72 and 98 DAA, and immediately stored in a Zip-top bag and stored in a Styrofoam box for transport to the laboratory to be used for the following analysis.

### Cell sap extraction using a cell pressure probe

The modified CPP technique (Hüsken et al. 1978) was used to collect cellular fluid from the target cells using a glass microcapillary. During the CPP operation, the oil pressure (turgor pressure when the tip was in the cell) and the meniscus location were measured using an automated cell pressure probe (ACPP, Wong et al. 2016) at 7.5 Hz, according to Wada et al. (2014). Microcapillary tips with long shanks were prepared by a Koph 750 micropipette puller and were beveled (Shackel et al. 1987) to give 2.5-4 μm i.d. tips utilizing a tip fabrication technique (Wada et al. 2011). All microcapillaries were autoclaved for 1 h at 180 °C. The tip was then set on the capillary holder and rinsed with a jet of DEPC-treated water twice and dried with an air jet.

The first extraction method, referred to as ‘shallow probing’, was conducted by penetrating skin cells. Prior to penetration, the oil pressure was set at 0.3 to 0.4 MPa, which is higher than turgor pressure (typically <0.2MPa) in skin cells at this developmental stage. The micropipette then penetrated the skin to a depth of between 20 μm and 230 μm below the epidermal surface. After reaching the target depth, oil pressure was slowly reduced until a meniscus could be observed in the microcapillary. The tip was then advanced until a rapid backward movement of the meniscus was observed, indicating penetration into a cell. Cell fluids were collected from at least two cells and the oil pressure was then reduced to −0.02 MPa and maintained. In some cases, fluids from the subepidermal cells located at 20 μm were collected by simply penetrating to this depth with an oil pressure of −0.02 MPa.

The second extraction method, referred to as ‘deep probing’, was used to obtain the fluid only from the mesocarp cells located from 1000 to 1500 μm below epidermis. To avoid possible contamination from the non-target (skin) cells, the oil pressure was pre-pressurized 0.3 to 0.4 MPa. The tip was then advanced to 1000 μm with a speed of of 75 μm/s. After reaching 1000 μm below epidermis, the pressure was slowly reduced to 0.03 MPa. The tip was then advanced, gradually reducing the oil pressure to −0.02 MPa in order to observe a rapid backward movement of the meniscus during the forward advance of the probe, typically collecting cellular fluids from two cells.

The third method, referred to as ‘continuous probing’ was used as a reference method to obtain the fluid from all cell layers. The oil pressure was maintained at −0.02 MPa, and the tip was advanced to between 1000 μm and 1350 μm from the epidermis, corresponding typically to 15-18 cell layers, with fluids collected throughout this range.

In all cases, the tip was quickly removed from the berry and submerged into a 10 μL droplet of DEPC-treated water on the inner wall of a 1.5mL autoclaved microcentrifuge tube. Applying a positive oil pressure, the fluid was quickly injected into the droplet and stored in a −80 °C until the further analysis. Additionally, DPEC water with no cell sap was used as negative control. In all cases, microcapillary tips were only used for one sample. The location of the meniscus formed in the microcapillary was tracked using the ACPP to determine the volume of cell sap injected. The range of the fluid volume collected by shallow, deep, and continuous probing was between 0.1-2.15 pL, 0.8-12.0 pL, and 5.0-12.0 pL, respectively.

### Gene expression analysis

Samples collected with the above extractions were centrifuged (to collect liquid at the bottom of the tube) and further used to quantify the expression of the UDP-glucose: flavonoid 3-O-glucosyltransferase (*VviUFGT*) gene, that codifies for the enzyme that determines the anthocyanin accumulation in the grape berry. *VviUFGT* is expressed in skin cells after veraison concomitantly with anthocyanin biosynthesis and accumulation in the vacuoles (Castellarin et al. 2007b, 2016).

Samples were divided in half as illustrated in **Fig. 2**. The first 5 μL were used for the expression assay of a housekeeping gene, *VvUbiquitin* (*VviUbi)* that is constitutively express in the cells, and another 5 μL were used for the expression assay of target gene, *VviUFGT.* Each qPCR reaction (10 μL) contained 250 nM in final of forward and reverse primers, 5 μL of sample, and 5 μL of Power SYBR^®^ green RNA-to-CT™ 1-step kit (Applied Biosystems) that has limits of detection as low as 2 copies of the target gene. The thermal cycling conditions were 95 °C for 10 min, followed by 95 °C for 15 s and 60 °C for 1 min for 40 cycles, according to the manual instruction. qPCRs were carried out as described in Castellarin et al. (2016). Primers pairs for *VviUFGT* and *VviUFGT* were designed on two exons flanking the intron. The expression of *VviUFGT* was analysed by these primer pairs: VviUFGT-forward (5’-GCAGGGCCTAACTCACTCTC-3’), VviUFGT-reverse (5’-AAATTGAGCAGCTCGTCTTCA-3’), and *VviUbi* was analyzed according to (Castellarin et al. 2007a). The final PCR products obtained from the *VviUFGT* amplification were analyzed in 1.5 % agarose gel. The samples that amplified two bands, 178bp (*VviUFGT* gDNA) and 101bp (*VviUFGT* mRNA) bands, were discarded because of the gDNA comtamination. The samples with only a 101bp band were considered to be positive samples and included in the data analysis. *VviUFGT* gene expression is reported relative to *VviUbi*.

## Declarations

### Ethics approval and consent to participate

Not applicable

### Consent for publication

Not applicable

### Availability of data and material

Not applicable

### Competing interests

The authors declare that they have no competing interests

### Funding

This work was supported by American Vineyard Foundation and in in part JSPS KAKENHI Grant Number 16H02533.

### Authors’ contributions

HW, SDC, and GAG conceived and designed the research, performed the experiments, analyzed the data and wrote the manuscript. All authors read, commented and approved the manuscript.

## Acknowledgements

Perseverance furthers.

## References

Bogs J, Downey MO, Harvey JS, et al (2005) Proanthocyanidin Synthesis and Expression of Genes Encoding Leucoanthocyanidin Reductase and Anthocyanidin Reductase in Developing Grape Berries and Grapevine Leaves. Plant Physiol 139:652–663. doi: 10.1104/pp.105.064238

Brandt, Kehr, Walz, et al (1999) Technical Advance: A rapid method for detection of plant gene transcripts from single epidermal, mesophyll and companion cells of intact leaves. Plant J 20:245–250

Brandt S, Kloska S, Altmann T, Kehr J (2002) Using array hybridization to monitor gene expression at the single cell level. J Exp Bot 53:2315–2323. doi: 10.1093/jxb/erf093

Brandt SP (2005) Microgenomics: gene expression analysis at the tissue-specific and single-cell levels. J Exp Bot 56:495–505. doi: 10.1093/jxb/eri066

Castellarin SD, Gambetta GA, Wada H, et al (2011) Fruit ripening in Vitis vinifera: spatiotemporal relationships among turgor, sugar accumulation, and anthocyanin biosynthesis. J Exp Bot 62:4345–54. doi: 10.1093/jxb/err150

Castellarin SD, Gambetta GA, Wada H, et al (2016) Characterization of major ripening events during softening in grape: turgor, sugar accumulation, abscisic acid metabolism, colour development, and their relationship with growth. J Exp Bot 67:709–22. doi: 10.1093/jxb/erv483

Castellarin SD, Matthews M a, Di Gaspero G, Gambetta G a (2007a) Water deficits accelerate ripening and induce changes in gene expression regulating flavonoid biosynthesis in grape berries. Planta 227:101–112. doi: 10.1007/s00425-007-0598-8

Castellarin SD, Pfeiffer A, Sivilotti P, et al (2007b) Transcriptional regulation of anthocyanin biosynthesis in ripening fruits of grapevine under seasonal water deficit. Plant Cell Environ 30:1381–1399. doi: 10.1111/j.1365-3040.2007.01716.x

Diakou P, Carde JP (2001) In situ fixation of grape berries. Protoplasma 218:225–35

Drea S, Derbyshire P, Koumproglou R, et al (2009) In situ Analysis of Gene Expression in Plants. In: Methods in molecular biology (Clifton, N.J.). pp 229–242

Efroni I, Birnbaum KD (2016) The potential of single-cell profiling in plants. Genome Biol 17:65. doi: 10.1186/s13059-016-0931-2

Gallagher JA, Koroleva OA, Tomos DA, et al (2001) Single cell analysis technique for comparison of specific mRNA abundance in plant cells. J Plant Physiol 158:1089–1092. doi: 10.1078/0176-1617-00285

Hüsken D, Steudle E, Zimmermann U (1978) Pressure probe technique for measuring water relations of cells in higher plants. Plant Physiol 61:158–63. doi: 10.1104/pp.61.2.158

Jones MA, Grierson CS (2003) A simple method for obtaining cell-specific cDNA from small numbers of growing root-hair cells in Arabidopsis thaliana. J Exp Bot 54:1373–8. doi: 10.1093/jxb/erg157

Kehr J (2003) Single cell technology. Curr Opin Plant Biol 6:617–21

Koroleva OA, Farrar JF, Tomos AD, Pollock CJ (1997) Patterns of Solute in Individual Mesophyll, Bundle Sheath and Epidermal Cells of Barley Leaves Induced to Accumulate Carbohydrate. New Phytol. 136:97–104

Laval V, Koroleva OA, Murphy E, et al (2002) Distribution of actin gene isoforms in the Arabidopsis leaf measured in microsamples from intact individual cells. Planta 215:287–92. doi: 10.1007/s00425-001-0732-y

Nakashima T, Wada H, Morita S, et al (2016) Single-Cell Metabolite Profiling of Stalk and Glandular Cells of Intact Trichomes with Internal Electrode Capillary Pressure Probe Electrospray Ionization Mass Spectrometry. Anal Chem 88:3049–57. doi: 10.1021/acs.analchem.5b03366

Nonami H, Schulze ED (1989) Cell water potential, osmotic potential, and turgor in the epidermis and mesophyll of transpiring leaves: Combined measurements with the cell pressure probe and nanoliter osmometer. Planta 177:35–46. doi: 10.1007/BF00392152

Schliep M, Ebert B, Simon-Rosin U, et al (2010) Quantitative expression analysis of selected transcription factors in pavement, basal and trichome cells of mature leaves from Arabidopsis thaliana. Protoplasma 241:29–36. doi: 10.1007/s00709-009-0099-7

Shackel KA, Matthews MA, Morrison JC (1987) Dynamic Relation between Expansion and Cellular Turgor in Growing Grape (Vitis vinifera L.) Leaves. Plant Physiol 84:1166–71. doi: 10.1104/pp.84.4.1166

Steudle E (1993) Pressure probe techniques: Basic principles and application to studies of water and solute relations at the cell, tissue and organ level. In: J S, H G (eds) Water deficits: Plant responses from cell to community. Bios Scientific Publishers, Oxford, pp 5–36

Tomos AD, Leigh RA (1999) The Pressure Probe: A Versatile Tool in Plant Cell Physiology. Annu Rev Plant Physiol Plant Mol Biol 50:447–472. doi: 10.1146/annurev.arplant.50.1.447

Wada H, Fei J, Knipfer T, et al (2014) Polarity of water transport across epidermal cell membranes in Tradescantia virginiana. Plant Physiol 164:1800–9. doi: 10.1104/pp.113.231688

Wada H, Hatakeyama Y, Onda Y, et al (2019) Multiple strategies for heat adaptation to prevent chalkiness in the rice endosperm. J Exp Bot 70:1299–1311. doi: 10.1093/jxb/ery427

Wada H, Matthews M a., Choat B, Shackel KA (2011) In situ Turgor Stability in Grape Mesocarp Cells and Its Relation to Cell Dimensions and Microcapillary Tip Size and Geometry. Environ Control Biol 49:61–73. doi: 10.2525/ecb.49.61

Wong DCJ, Lopez Gutierrez R, Dimopoulos N, et al (2016) Combined physiological, transcriptome, and cis-regulatory element analyses indicate that key aspects of ripening, metabolism, and transcriptional program in grapes (Vitis vinifera L.) are differentially modulated accordingly to fruit size. BMC Genomics 17:416. doi: 10.1186/s12864-016-2660-z

Yuan G-C, Cai L, Elowitz M, et al (2017) Challenges and emerging directions in single-cell analysis. Genome Biol 18:84. doi: 10.1186/s13059-017-1218-y

